# Intermittent self-administration of fentanyl induces a multifaceted addiction state associated with persistent changes in the orexin system

**DOI:** 10.1101/2020.04.23.055848

**Authors:** Jennifer E. Fragale, Morgan H. James, Gary Aston-Jones

## Abstract

The orexin (hypocretin) system plays a critical role in motivated drug-taking. Cocaine self-administration with the intermittent access (IntA) procedure produces a robust addiction-like state that is orexin-dependent. Here, we sought to determine the role of the orexin system in opioid addiction using IntA self-administration of fentanyl. Different groups of male rats were either given continuous access in 1h (short access; ShA), or 6h periods (long access, LgA), or IntA (5min of access separated by 25min of no-access) to fentanyl for 14 days. IntA produced a greater escalation of fentanyl intake, motivation for fentanyl on a behavioral economics task, persistent drug seeking during abstinence, and cued-induced reinstatement compared to rats given ShA or LgA. We found that addiction behaviors induced by IntA to fentanyl were reversed by the orexin-1 receptor antagonist SB-334867. IntA to fentanyl was also associated with a persistent increase in the number of orexin-expressing neurons. Together, results indicate that the IntA model is a useful tool in the study of opioid addiction, and that the orexin system is critical for the maintenance of addiction behaviors induced by IntA self-administration of fentanyl.

## Introduction

Orexins (hypocretins) are neuropeptides exclusively produced in the hypothalamus that regulate a wide range of behaviors, including feeding, arousal, and drug reward (de Lecea et al., 1998; Harris et al., 2005; Sakurai et al., 1998^1^. Increased orexin cell number and activity occur in rats that exhibit enhanced ‘addiction-like’ states^2–5^. We found that rats with a strong, addiction-like state for cocaine (e.g., following intermittent access, IntA; described below) persistently exhibit greater numbers of lateral hypothalamic (LH) orexin-expressing neurons than animals with a milder addiction profile^5^. We also reported that the number of LH orexin-expressing neurons predicts baseline (pre-IntA) motivation for cocaine^3^. Recent studies also revealed that higher numbers of orexin neurons are found in postmortem brains of heroin addicts than in non-addict control brains^6^. That study also reported that the number of orexin-expressing neurons increased in mice following non-contingent (experimenter administered) injections of morphine.

Orexins regulate drug-seeking behaviors primarily through signaling at orexin-1 receptors (Ox1Rs)^7–15^. OxR1 antagonists are particularly effective in attenuating addiction behaviors during enhanced ‘addiction-like’ states, as the selective Ox1R antagonist SB-334867 (SB) is most effective at increasing economic demand elasticity (decreasing motivation) for cocaine in rats with high baseline demand^16^. In addition, SB only reduces cocaine intake in animals following extended drug access^17^. Moreover, rats that exhibit stronger addiction-like behaviors (e.g., following IntA) are more sensitive to OxR1 antagonism such that lower doses of SB attenuate addiction behaviors than in animals given short or long access to drug^5^. Our lab recently reported similar results for opioids: Systemic administration of SB preferentially attenuated economic demand for fentanyl in rats with high baseline demand^9^, and local microinjections of SB into ventral pallidum (VP) produced identical results for remifentanil^15^.

The IntA paradigm is argued to better recapitulate patterns of drug intake in clinical populations compared to more commonly used paradigms in which drug is continuously available^18,19^. In the case of cocaine, IntA produces a multifaceted animal model of addiction that includes robust escalation of intake, increased motivation for cocaine in a behavioral economics (BE) task, increased compulsive drug taking, and greater reinstatement of drug seeking compared to traditional short (ShA) or long (LgA) continuous access models^5,20–24^. Although the IntA model was recently applied to heroin self-administration^25^, the utility of the IntA paradigm to induce an addiction multiphenotype for opioids has not been directly compared with the more widely-utilized long access (LgA) model^26^.

Here, we employed IntA to the widely abused opioid fentanyl, and directly compared addiction-like behaviors to those for the LgA model of addiction. We found that IntA self-administration of fentanyl is associated with development of a multifaceted addiction-like phenotype characterized by escalation of intake, increased economic demand, persistent drug seeking during abstinence, and more significant reinstatement of drug seeking compared to rats given LgA or short access (ShA) to fentanyl. We also found that IntA to fentanyl is associated with higher numbers of orexin-expressing neurons, and that addiction behaviors in IntA rats are more sensitive to Ox1R antagonism compared to ShA rats. Together, these findings indicate that IntA self-administration provides a strong model of opioid addiction, and that augmentation of the orexin system likely contributes to the corresponding enhanced addiction-like state.

## Methods

### Subjects

Sprague Dawley male rats (250-300g) were obtained from Charles River Laboratories (Kingston, NY). Rats were pair-housed under a 12:12 h reverse light cycle (lights off at 0800h and on at 2000h) and given *ad libitum* access to food and water. All procedures were conducted in accordance with the NIH Guide for the Care and Use of Laboratory Animals and approved by Rutgers University New Brunswick Institutional Animal Care and Use Committee.

### Drugs

Fentanyl HCl powder and the selective OxR1 antagonist SB-334867 (SB) were obtained from the National Institute of Drug Abuse Drug Supply Program. Fentanyl HCl powder was dissolved in 0.9% sterile saline to a concentration of 8μg/ml. SB was prepared as previously described and injected i.p. at a volume of 4.0 ml/kg^8,12,13,15^.

### Catheter Implantation

Chronic indwelling jugular vein catheters were implanted as previously described^14^. Rats were anesthetized with 2% isoflurane and administered the analgesic rimadyl (5 mg/kg, s.c.) prior to catheter implantation. After surgery, catheters were flushed daily with the antibiotic cefazolin (0.1 mL; 100mg/mL) and heparin (0.1 ml; 100 U/ml). Rats were allowed one week to recover prior to self-administration training.

### Self-Administration Training

Rats were trained in Med Associates operant boxes housed in individual sound-attenuating chambers (Med Associates, St Albans, VT, USA). Operant boxes were fitted with 2 levers (active and inactive), a stimulus light located directly above the active lever, speaker, and house light. All boxes were controlled by Med-PC IV software (Med Associates, St Albans, VT, USA). Rats were first trained in 2 hr sessions on a fixed-ratio 1 (FR-1) schedule. Active lever presses resulted in a 3.6s infusion of 0.5μg of fentanyl paired with stimulus light (white light) and tone (78 dB, 2900 Hz). Each infusion was proceeded by a 20-sec time out signaled by termination of the house light. Rats were trained for a minimum of 6 sessions and to a criterion of > 25 infusion for 3 consecutive sessions.

### Baseline Fentanyl Demand Testing

Fentanyl demand was assessed using a within-session BE procedure as previously described^9^. Briefly, animals were tested in 110-minute sessions where the dose of fentanyl per infusion was decreased on a quarter logarithmic scale in successive 10-minute bins. Demand curves were generated using the exponential demand equation^27^ and a least sum of squared differences procedure for curve fitting^28,29^. All data points up until two bins past the point of maximal price paid per μg fentanyl (P_max_) were included in the analysis. Demand curves were generated for individual subjects for individual BE sessions and demand parameters Q_0_ and alpha (a) were calculated. Q_0_ is a theoretical measure of drug consumption when no effort is required and is also a measure of a subject’s hedonic set point. a is a measure of demand elasticity (price sensitivity of consumption) and is used as an inverse measure of motivation. Rats underwent a minimum of 6 BE sessions and until Q_0_ and a values differed by less than 25% across three consecutive sessions.

### Short, Long and Intermittent Access Groups

To ensure baseline a values did not differ between access groups, rats were pseudorandomly assigned to short access (ShA), long access (LgA) or intermittent (IntA) self-administration following initial assessment of fentanyl demand. All access groups were given their access treatment for 14 consecutive daily sessions. ShA rats were given continuous access to fentanyl on an FR1 schedule in 1h sessions. LgA rats received continuous access to fentanyl on an FR1 schedule during 6h sessions. In both procedures, active lever responses resulted in a 3.6s infusion of 0.5μg of fentanyl paired with light and tone. Each infusion was proceeded by a 20s time-out period signaled by termination of the house light. For IntA rats, access was similar to that previously described for cocaine^5,21^. Briefly, fentanyl was available in 5min bins separated by 25min periods of non-availability. Drug availability periods were signaled by a priming infusion of fentanyl (1s infusion of 0.14μg of fentanyl) paired with light and tone. Active lever presses made during this time resulted in a 3.6s infusion of 0.5μg of fentanyl paired with light and tone. Unlike ShA and LgA, during IntA infusions were not proceeded by a 20s time-out period. After 5 min, levers were retracted, and the house light was terminated.

### Post-Access Fentanyl Demand and Pharmacological Testing

One day following the final ShA, LgA or IntA session, rats were again tested for fentanyl demand using the BE procedure described above. Demand was measured over a minimum of 6d and until stable demand values were observed over 3 consecutive days. In rats receiving pharmacological testing, SB (0, 10 or 30 mg/kg) pretreatment was given 30min prior to testing in a within-subjects, counterbalanced design with a minimum of 3 d between tests.

### Extinction Training and Cued-Induced Reinstatement

Following BE testing, ShA, LgA, and IntA rats were given 2h extinction sessions during which active lever presses were no longer paired with infusions or fentanyl-associated light/tone cues. Extinction sessions continued for a minimum of 7 d and until active lever presses in the final 3 sessions were ≤ 25 presses. The following day, rats were tested for cued reinstatement of fentanyl seeking, whereby active lever presses produced fentanyl-associated cues (light and tone). In rats receiving pharmacological testing, SB pretreatment (0, 10 or 30 mg/kg) was given 30min prior to testing in a within-subjects, counterbalanced design with a minimum of 3 d between reinstatement tests to avoid any potential carryover effects of treatment^15^.

### Tissue Preparation for Immunohistochemistry

Ninety days after their final BE test, rats were deeply anesthetized with sodium pentobarbital and transcardially perfused with 0.9% sterile saline then 4% paraformaldehyde. Brains were harvested and postfixed overnight in 4% paraformaldehyde. The next day, brains were transferred to a 20% sucrose-PBS azide solution and stored at 4°C until sectioned. Brains were sectioned at 40μm using a cryostat and sections were stored in PBS azide at 4°C.

### Immunohistochemistry

Tissue was incubated in mouse anti-orexin A (1:500, Santa Cruz Biotechnology, catalog number sc-80263, JCN Antibody Database AB_1126868) and rabbit anti-MCH (1:5000, Phoenix Pharm, catalog number H-070-47, JCN Antibody Database AB_10013632) antibodies in 5% normal donkey serum overnight at room temperature. The following day, tissue was incubated in Alexa-Fluor 594 conjugated donkey anti-mouse and Alexa-Fluor 488 conjugated donkey anti-rabbit for 2h at room temperature. Sections were rinsed in phosphate buffered saline, mounted onto glass slides and cover slipped using Fluoroshield Mounting Medium with DAPI (Abcam).

### Cell Quantification

Quantification of orexin- and MCH-expressing neurons was performed as previously described^3,5,30,31^. Briefly, coronal images of the orexin cell field (2.5-3.8 mm caudal to bregma) were taken using a Zeiss Axio Zoom V16 microscope. Tiled photographs were compiled at 20x magnification using Zen 2 imaging software (Carl Zeiss Microscopy). Counts of orexin A- or MCH-expressing neurons were taken in both hemispheres using ImageJ software (NIH) by two investigators blinded to the experimental condition. Medial and lateral hypothalamic subregions of the orexin cell field were quantified separately due to evidence of a functional dichotomy between these populations^32^. Medial and lateral cell populations were divided by a vertical line made 100μm lateral to the fornix. As in our previous studies, three sections per animal were analyzed and the average of neurons counted in both hemispheres in each section were determined and averaged across the three sections^3,5^.

### Data Analysis

Data were expressed as mean values ± 1 standard error of the mean. Statistics were performed using GraphPad Prism for Mac (Version 7, GraphPad Software Inc., La Jolla, CA) with an α level of 0.05. First-hour escalation of fentanyl intake was assessed using separate repeated-measures ANOVAs with Dunnett’s multiple comparisons as *post hoc* tests. Demand parameters Q_0_ and a were expressed as percent change from pre-access baseline and analyzed using separate repeated-measures ANOVAs. Dunnett’s multiple comparisons test were used as *post hoc* tests when allowed. All between access group comparisons were made using one-way ANOVAs with Sidak’s *post hoc* test. Extinction and reinstatement data were assessed using mixed-design ANOVAs with Sidak’s *post hoc* tests. A Kaplan-Meier estimator was used to compare extinction rates between access groups. Independent samples t-tests were used to analyze orexin and MCH cell counts between ShA and IntA rats.

## Results

### IntA to Fentanyl is Associated with a Multifaceted ‘Addiction-Like’State

Figure 1A outlines the experimental timeline. Rats were randomly assigned to ShA, LgA or IntA following baseline BE testing. Baseline (pre-access treatment) a and Q_0_ values were similar between rats assigned to ShA, LgA and IntA (F_2,29_= 1.386, p=0.266, F_2,29_= 0.333, p=0.719, respectively, one-way ANOVAs; data not shown). Escalation of first-hour fentanyl intake was not observed in ShA rats (rm-ANOVA, F_13,143_= 1. 104, p=0.360) or LgA (rm-ANOVA, F_13,78_= 1.042, p=0.4213). In contrast, significant escalation of first-hour fentanyl intake was observed in IntA rats (Figure 1B; rm-ANOVA, F_13,156_= 3.645, p<0.001). Overall fentanyl consumption on day 14 of access treatment was greatest in LgA rats (Figure 1C; one-way ANOVA, F_2,28_= 13.79, p<0.0001 with Sidak’s test, LgA v. ShA, p<0.0001, and LgA v. IntA, p=0.005). Fentanyl consumption did not differ between ShA and IntA rats (ShA v. IntA, p=0.1191).

**Figure 1.**
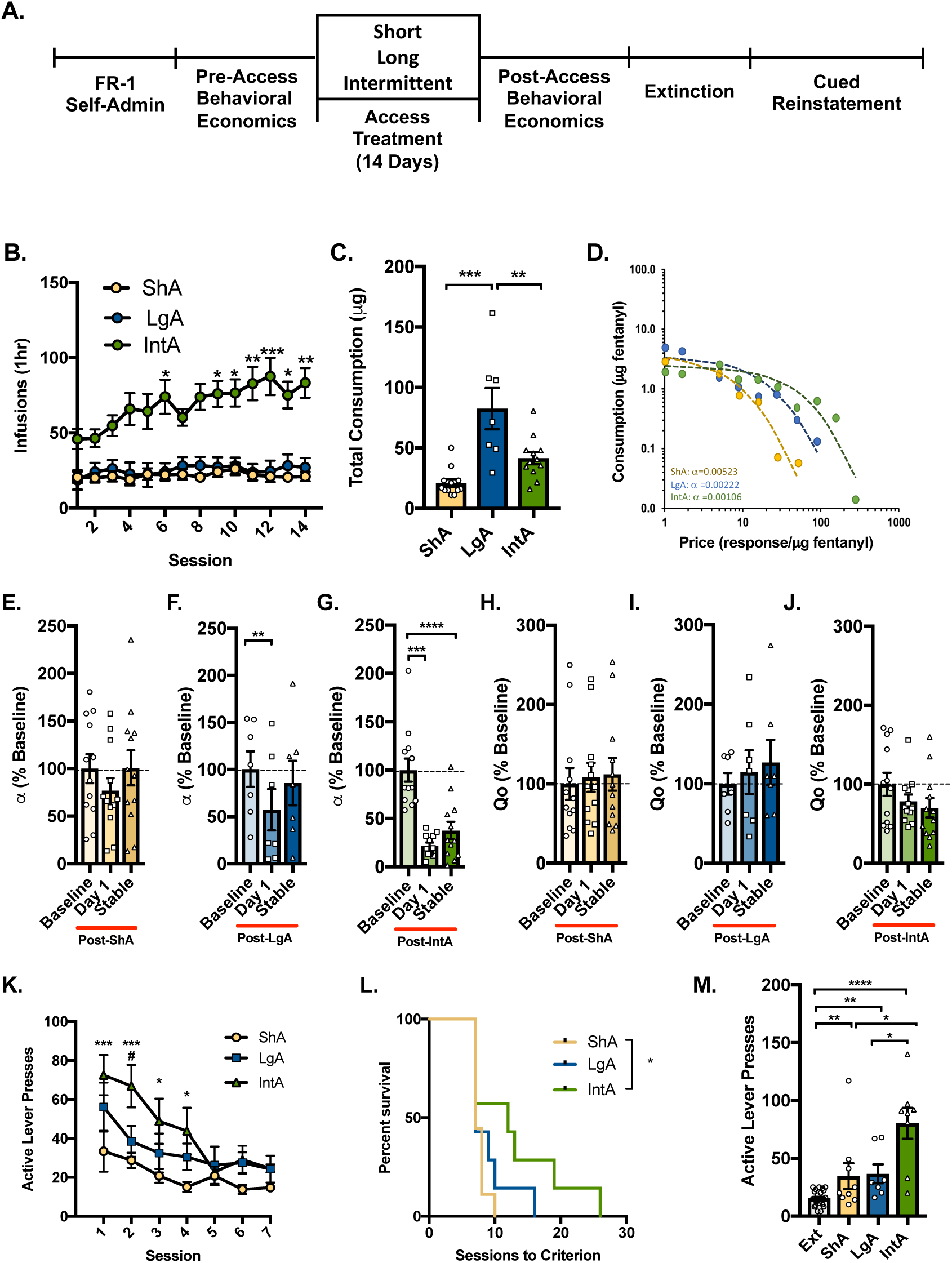
IntA to fentanyl promotes a strong ‘addiction-like’ state. A) Timeline of behavioral testing. Rats were assigned to short access (ShA; yellow; n=12), long access (LgA; blue; n=7), or intermittent access (IntA; green; n=12) following initial demand testing. B) First-hour fentanyl intake for different groups across the 14d of access treatment. Escalation of first-hour fentanyl intake was observed with IntA, but not ShA or LgA [*denotes significant difference compared to day 1; rm-ANOVA with Dunnett’s post hoc test]. C) Overall consumption of fentanyl was greater with LgA compared to ShA or IntA (data shown for day 14; one-way ANOVA with Sidak’s post hoc test]. There was no statistical difference in overall intake between ShA and IntA groups (p<*). D) Representative demand curves from single subjects after ShA, LgA, or IntA to fentanyl. E) Motivation (a) did not differ from pre-access baseline following ShA to fentanyl. F) LgA to fentanyl was associated with a transient decrease in a (increased motivation) on the first day but not when BE performance stabilized (∼1-week post LgA) [rm-ANOVA with Dunnett’s post hoc test]. G) IntA was associated with a robust decrease in a (increased motivation) on the first day and after BE performance stabilized (∼1-week post-IntA) compared to pre-access baseline [rm-ANOVA with Dunnett’s post hoc test]. H-J) ShA, LgA, and IntA to fentanyl had no significant effect on the free consumption of fentanyl (Q_0_) [rm-ANOVA with Dunnett’s post hoc test]. K) Following BE testing, lever press responding was extinguished in ShA (n=9), LgA (n=7), and IntA (n=8) rats. Rats given IntA to fentanyl made more active lever presses during the first four extinction sessions compared to ShA. Active lever presses differed to a lesser extend between IntA and LgA rats and did not differ between ShA and LgA rats [mixed-design ANOVA with Sidak’s post hoc test; * ShA v. IntA; # LgA v. IntA]. L) Rats given IntA to fentanyl also required more extinction sessions to reach extinction criterion compared to ShA rats. However, sessions to extinction criterion were similar between LgA and IntA rats (Kaplan-Meier estimator). M) All access groups showed significant reinstatement of drug seeking (active lever pressing) in response to fentanyl associated cues [mixed-design ANOVA with Sidak’s post hoc test]. However, cue-induced responding was greater in rats given IntA compared to rats given ShA or LgA [extinction data pooled; one-way ANOVA with Sidak’s post hoc test]. For all panels, # p<0.05, *p<0.05, **p<0.01, ***p<0.001, ****p<0.0001.

Demand for fentanyl was reassessed in rats following ShA, LgA, or IntA. Figure 1D shows representative demand curves from individual ShA (yellow), LgA (blue), and IntA (green) subjects. Motivation for fentanyl (inverse a) was stable during ShA and did not differ from pre-access baseline (Figure 1E; rm-ANOVA, F_2, 22_ = 1.443, p=0.2576). LgA produced only a transient increase in motivation (decreased a on Day 1) that dissipated in < ∼7d post-LgA (Figure 1F; rm-ANOVA, F_2,14_ = 7.558 p=0.0059 with Dunnett’s multiple comparisons test, Baseline v. Day 1, p<0.0018 and Baseline v. Stable, p=0.276). In contrast, IntA was associated with a robust and persistent increase in motivation for fentanyl (decreased a), such that a values on the first BE session following IntA were significantly lower compared to baseline a (Figure 1G; rm-ANOVA, F_2, 22_ = 31.86, p<0.0001 with Dunnett’s multiple comparisons test, Baseline v. Day 1 p<0.0001), and persisted for > ∼7d following IntA (Baseline v. Stable p<0.0001). No relationship was observed between fentanyl intake and changes in a in rats previously IntA to fentanyl (Pearson’s r, R^2^=0.0603, p=0.4417; data not shown). ShA, LgA and IntA to fentanyl were not associated with changes in Q_0_ (Figures 1H-1J; F_2, 22_ = 1.362, p=0.2270; F_2, 14_ = 0.6434, p=0.5404; F_2, 22_ = 2.981, p=0.0715, respectively, rm-ANOVAs).

### IntA Produces Greater Extinction Responding

Following BE testing, ShA, LgA, and IntA rats underwent extinction training where active lever presses produced neither fentanyl nor fentanyl-associated cues. Overall rates of active lever presses decreased for all access groups over the first 7 extinction sessions (Figure 1K; mixed-design ANOVA; main effect of Session F_6, 126_ = 13.09, p<0.0001). Active lever responding differed among access groups (main effect of Condition F_2,21_ = 4.923, p<0.0001 and Condition x Session interaction F_12,126_ = 1.84, p=0.0484). ShA and LgA rats did not differ in the number of active lever presses made across extinction sessions (Sidak’s test p>0.05). IntA rats made more active lever presses across the first 4 extinction sessions compared to rats given ShA (Sidak’s test p>0.05). Active lever presses differed between IntA and LgA rats to a lesser extent (Second extinction session; Sidak’s test, p>0.05). Days to extinction criterion also differed among access groups (Figure 1L; Kaplan-Meier estimator, χ^2^_(1)_=5.252, p=0.0219). IntA rats required more extinction sessions to reach extinction criterion compared to rats given ShA (Kaplan-Meier estimator, χ^2^_(1)_=4.491, p=0.0341. Sessions to extinction criterion did not differ significantly between rats given ShA vs. LgA, or between those given LgA vs. IntA to fentanyl (Kaplan-Meier estimator, χ^2^_(1)_=1.324, p=0.250; χ^2^_(1)_=1.931, p=0.164). Median sessions to extinction criterion (± SEM) were 7.0 ± 0.3 for ShA, 9.0 ± 1.2 for LgA, and 11.8 ± 2.2 for IntA.

### IntA Produces Greater Reinstatement Responding

After reaching extinction criterion, rats were tested for cued reinstatement of fentanyl seeking. Here, active lever responses produced fentanyl-associated light+tone compound cues but no drug delivery. In all access groups, fentanyl-associated cues induced significant reinstatement of drug seeking as measured by increased responding on the active lever (mixed-design ANOVA; ShA: time x lever Interaction F_1,16_=4.624, p=0.0474 with Sidak’s test p=0.0282; LgA: time x lever Interaction F_1,12_=7.808, p=0.0162 with Sidak’s test p=0.0054; IntA: time x lever Interaction F_1,14_=22.17, p=0.0003 with Sidak’s test p<0.0001). Reinstatement of drug seeking differed among access groups (Figure 1M; one-way ANOVA, F_2,21_= 5.186, p=0.0148). ShA and LgA rats did not differ in reinstatement responding (Sidak’s test, ShA v. LgA, p=0.9916). In contrast, rats given IntA to fentanyl made more active lever presses in response to fentanyl-associated cues compared to ShA or LgA rats (Sidak’s test, ShA v. IntA, p=0.0241, and LgA v. IntA, p=0.0468).

### Ox1R Signaling Underlies the Maintenance of IntA-Induced ‘Addiction-Like’ Behaviors

To determine if Ox1R signaling was critical for the maintenance of IntA-induced ‘addiction-like’ behaviors, a subset of ShA and IntA rats received SB (0, 10, or 30 mg/kg, ip) prior to a BE test and cue-induced reinstatement. Figure 2A outlines the experimental timeline. In these experiments, ShA rats were used as the comparison group for IntA because total fentanyl consumption was similar between ShA and IntA rats (described above). InShA rats, only 30 mg/kg SB was effective in reducing motivation (increasing a; Figure 2B; rm-ANOVA, F_3, 24_ = 8.922, p=0.0004, Dunnett’s multiple comparison test Vehicle v. SB 30, p=0.0006). SB significantly reduced motivation for fentanyl in rats given IntA (increased a; Figure 2C; rm-ANOVA, F_3, 18_= 9.179, p=0.0007). *Post hoc* analyses revealed that pretreatment with either 10 mg/kg or 30mg/kg SB decreased motivation (increased a) relative to vehicle pretreatment (Dunnett’s multiple comparison test Vehicle v. SB 10, p=0.0093, Vehicle v. SB 30, p=0.0008). Vehicle pretreatment had no effect on IntA-induced increases in motivation (Pre-IntA Baseline v. Vehicle, p=0.0007).

**Figure 2.**
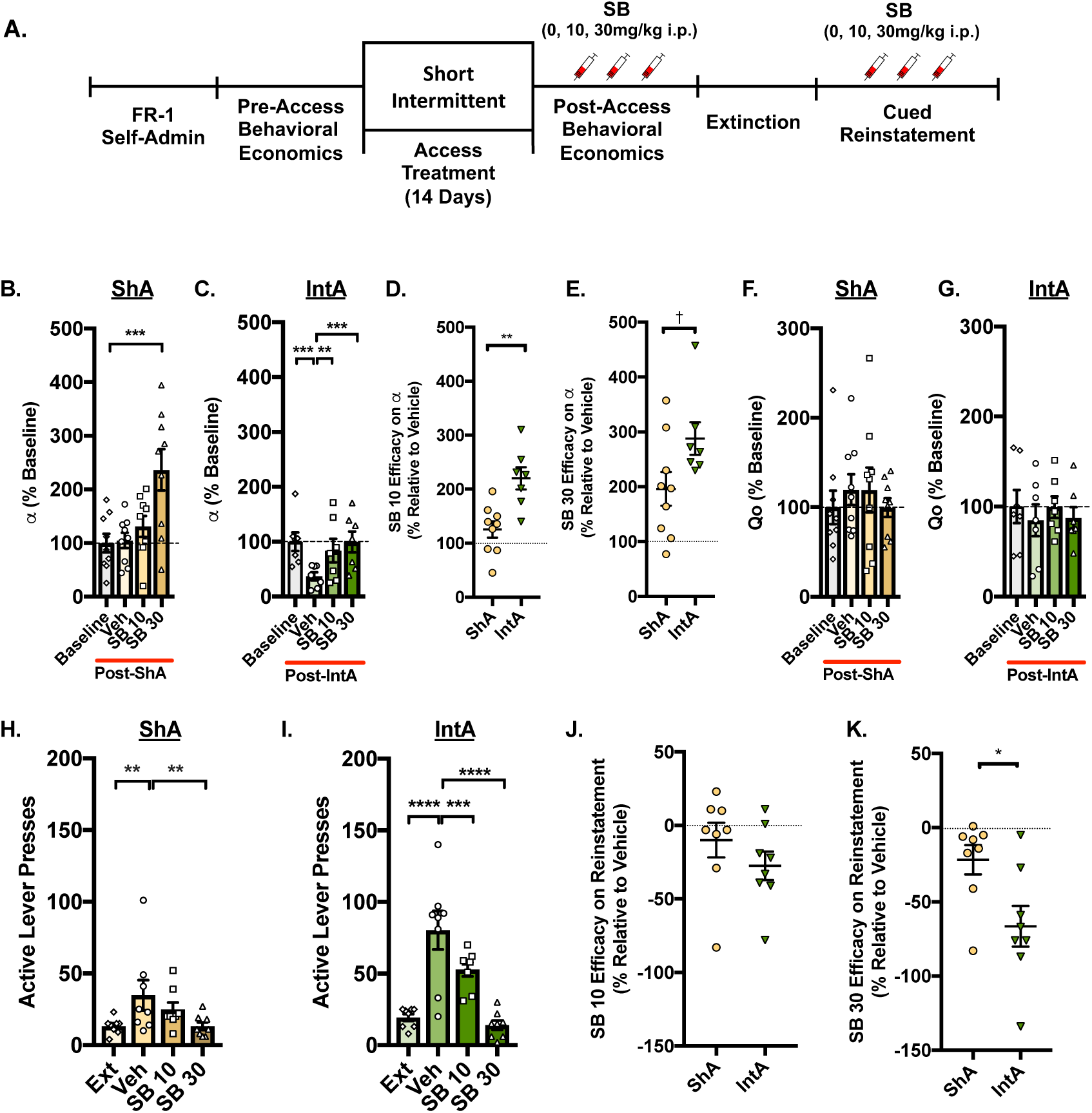
Ox1R signaling promotes the IntA-induced ‘addiction-like’ state. A) Timeline of behavioral and pharmacological testing. Prior to BE testing, the Ox1R antagonist SB (10 or 30mg/kg, ip) or vehicle was given to a subset of ShA (n=9) and IntA (n=7) rats. B) In ShA rats, SB increased a (decreased motivation) relative to vehicle only at the highest dose tested [30mg/kg; rm-ANOVA with Dunnett’s post hoc test]. C) In contrast, in IntA animals 10 or 30mg/kg SB increased a values (decreased motivation) relative to vehicle [rm-ANOVA with Dunnett’s post hoc test]. D-E) 10 or 30mg/kg SB was more effective at reducing motivation (increasing a) in rats previously given IntA to fentanyl compared to rats given ShA. F-G) SB had no effect on Q_0_ in ShA or IntA rats. H) In ShA rats (n=8), fentanyl-associated cues reinstated active lever pressing after extinction of fentanyl self-administration, but only SB 30mg/kg reduced this responding [mixed-design ANOVA with Sidak’s post hoc]. I) The presentation of fentanyl-associated cues also reinstated active lever pressing following extinction in rats given IntA (n=8). Unlike ShA rats, 10 or 30mg/kg SB attenuated this effect [mixed-design ANOVA with Sidak’s post hoc]. J) Low dose SB (10mg/kg) was similarly effective in attenuating cued reinstatement (reducing active lever presses) in ShA and IntA rats. K) In contrast, pretreatment with SB (30 mg/kg) was more effective in attenuating cued reinstatement in IntA rats. †=0.0544, *p<0.05, **p<0.01, ***p<0.001, ****p<0.0001.

The extent to which SB affected motivation (SB efficacy) was compared for different doses in rats given ShA vs. IntA. SB at 10mg/kg was more effective in reducing motivation (increasing a) in rats given IntA compared to those given ShA (Figure 2D; t_(14)_=3.783, p=0.002). Similar results were seen for SB at 30mg/kg but failed to reach statistical significance (Figure 2E; t_(14)_=2.10, p=0.0544). SB had no effect on Q_0_ in rats given ShA or IntA to fentanyl (Figures 2F-G; F_2, 24_ = 0.9583, p=0.4305; F_3, 18_ = 0.6872, p=0.5717, respectively).

The effects of SB on cue-induced reinstatement responding were also assessed in rats given ShA or IntA to fentanyl. SB was effective in attenuating cue-induced reinstatement in ShA rats (Figure 2H; mixed-design ANOVA; main effect of Treatment F_1,14_=22.65, p<0.0003). In ShA rats, fentanyl-associated cues reinstated extinguished drug seeking (Sidak’s multiple comparison test Ext v. Vehicle, p=0.0053); however, SB was only effective in attenuating this behavior at 30mg/kg dose (Vehicle v. SB 30, p=0.0053). SB had no effect on inactive lever responding (Vehicle v. SB 10, p>0.999; Vehicle v. SB30, p=0.991, data not shown). In IntA rats, SB dose-dependently attenuated cue-induced reinstatement (Figure 2I; mixed-design ANOVA; main effect of treatment F_1,14_=60.82, p<0.0001; treatment x lever Interaction F_3,42_=21.63, p<0.0001). In these rats, the presentation of fentanyl-associated cues reinstated extinguished drug seeking (Sidak’s multiple comparison test Ext v. Vehicle, p<0.0001), and SB (10 and 30 mg/kg) significantly attenuated this behavior (Vehicle v. SB 10, p=0.0007; Vehicle v. SB 30, p<0.0001). SB had no effect on inactive lever responding (Vehicle v. SB 10, p>0.999; Vehicle v. SB30, p>0.999; data not shown).

As shown in Figures 2J, pretreatment with 10mg/kg SB was similarly effective at attenuating cue-induced reinstatement in rats previously given ShA and IntA to fentanyl (t_(14)_=1.147, p=0.2704). However, pretreatment with 30mg/kg SB was most effective at attenuating cue-induced reinstatement in rats previously given IntA to fentanyl (Figure 2K; t_(14)_=2.648, p=0.0191).

### Increased Numbers of Orexin-Expressing Neurons Following IntA to Fentanyl

Subgroups of ShA and IntA rats were sacrificed following 90d of abstinence to investigate the effect of differential fentanyl access on numbers of orexin- and MCH-expressing neurons. Figure 3A displays images of orexin- (top panel) and MCH-expressing neurons (middle panel) in the hypothalamus of an IntA rat. MCH-expressing neurons are unique to the hypothalamus and are present where orexin-expressing neurons are located (bottom panel)^33,34^. Overall, rats given IntA to fentanyl had a greater number of orexin-expressing neurons compared to rats given ShA (independent samples, t_(10)_=3.049, p=0.0123). Compared to ShA, rats given IntA to fentanyl had higher numbers orexin-expressing neurons in both the dorsomedial/perifornical hypothalamus (DMH/PF; Figure 3B) and lateral hypothalamus (LH; Figure 3C; independent samples, t_(10)_=2.848, p=0.0173; t_(10)_=2.585, p=0.0272, respectively). Differences in cell numbers were specific to orexin neurons, as the numbers of MCH-expressing neurons in the DMH/PF and LH did not differ between ShA and IntA rats (Figure 3D-E; t_(9)_=0.9713, p=0.3568; t_(9)_=1.465, p=0.1769, respectively).

**Figure 3.**
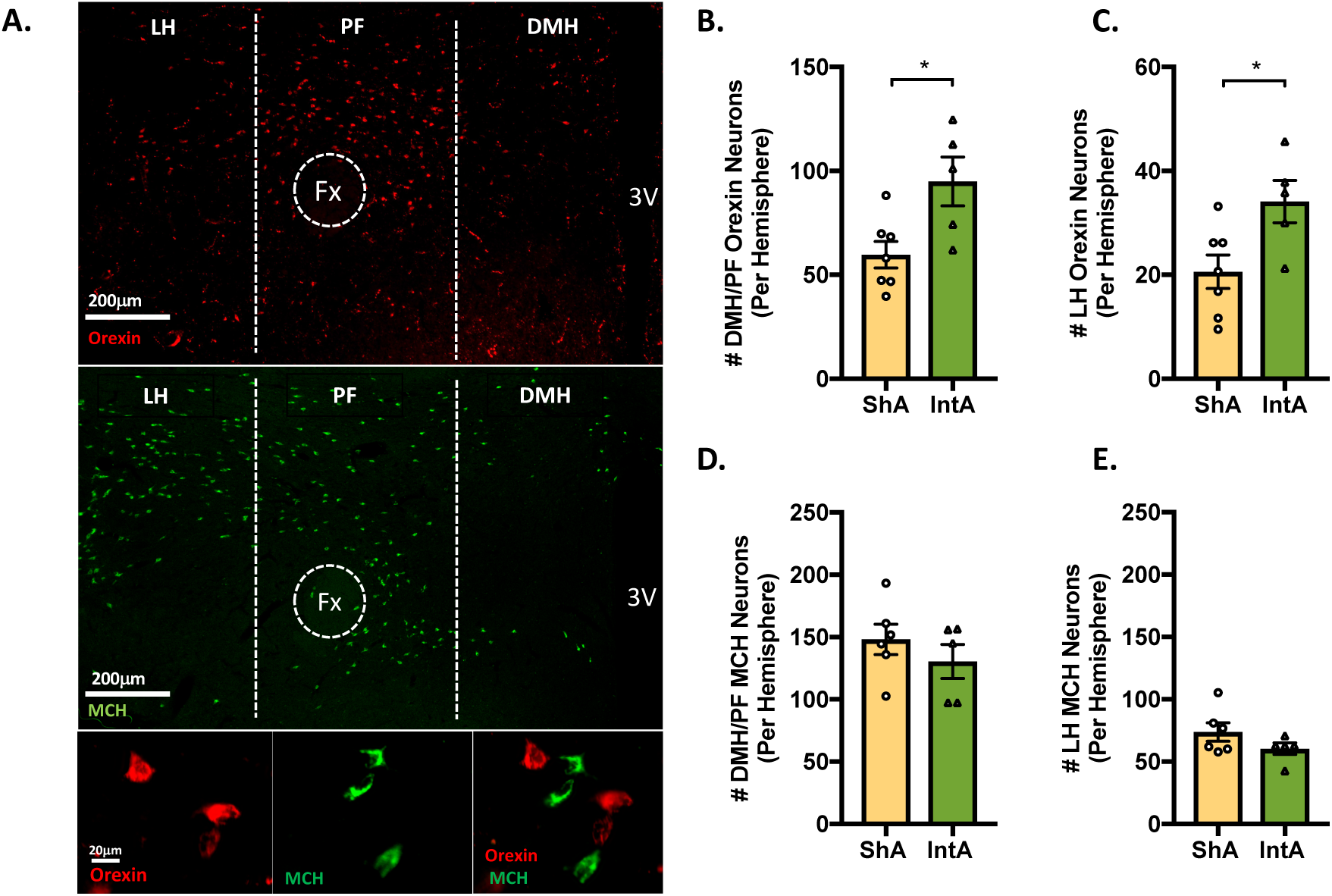
IntA to fentanyl is associated with increased number of orexin-expressing neurons. Orexin- and MCH-immunoreactive neuronal numbers were assessed after 90 days of abstinence in rats previously given ShA (n=7) or IntA (n=5) to fentanyl. A) Low magnification photomicrograph of a frontal section showing orexin (red; top panel) and MCH-expressing neurons (green; middle panel) in the hypothalamus of a rat given IntA to fentanyl. High-magnification photomicrograph showing orexin (red; bottom left), MCH (green; bottom middle), and merge (bottom right). B-C) Compared to ShA, rats given IntA to fentanyl had a greater number of orexin-expressing neurons in the both the DMH/PF and LH orexin cell fields [independent samples t-tests]. D-E) In contrast, the number of DMH/PF and LH MCH-expressing neurons did not differ between ShA and IntA rats. For all panels, *p<0.05

## Discussion

Here, we characterized the addiction phenotype induced by IntA to the opioid fentanyl, and directly compared these results to those for the more commonly utilized LgA or ShA models of addiction. We found that IntA to fentanyl produced greater escalation of fentanyl intake, increased motivation for fentanyl on a BE task, greater persistent drug seeking during abstinence/extinction, and higher cue-induced reinstatement of extinguished fentanyl seeking compared to LgA or ShA animals. In addition, we investigated the role of the orexin system in the maintenance of these behaviors. We found that the IntA-induced increases in fentanyl demand and reinstatement were reversed by pretreatment with the selective Ox1R antagonist SB at doses lower than what was required to produce similar effects in ShA rats. The efficacy of SB in attenuating these behaviors was also greater in IntA rats. Finally, we found that IntA was associated with a persistent increase in the number of orexin-expressing neurons. Together, these results demonstrated that the IntA procedure is a robust model by which to induce a multifaceted opioid addiction phenotype, and that the orexin system is critical for the maintenance of IntA-induced addiction behaviors.

### Intermittent Access as a Model for Opioid Addiction

Extended drug access (LgA; ∼6-12h/day) captures key elements of addiction behavior^26,35,36^. Rats given LgA to opioids such as heroin, oxycodone, or fentanyl showed dose-dependent escalation of drug intake and greater responding for drug on progressive ratio (PR) tasks compared to rats given ShA^37^. Consistent with these findings, we demonstrated that LgA to fentanyl was associated with increased motivation for fentanyl (lower a) relative to baseline. However, we also found that the increase in motivation following LgA to fentanyl was transient as a values returned to baseline ∼7 days post-LgA. Moreover, IntA but not LgA rats showed greater extinction responding and cue-induced reinstatement of fentanyl seeking compared to ShA rats. Our lab and others found that IntA to cocaine induced similarly enhanced addiction behaviors in rats, including increased motivation and compulsive responding for drug, and greater cued and drug-primed reinstatement of drug seeking compared to ShA or LgA rats^5,20,23^. Interestingly, a recent study reported individual differences in several addiction behaviors following IntA to heroin indicating that some rats might be particularly vulnerable to developing an addiction phenotype in response to IntA opioid self-administration^25^. Here, we show that at a population level, IntA offers a robust model by which to induce a multifaceted ‘addiction-like’ state for opioids.

In the present study, we did not observe escalation of fentanyl intake with LgA as seen with consumption in the first hour. Previous studies using a similar dose of fentanyl reported similar findings^37^. Importantly, we found no relationship between the degree of escalation during IntA access to fentanyl and the magnitude of change in motivation (a) for fentanyl, indicating that subsequent changes in motivation for fentanyl are not dependent on escalation of drug intake.

### Role of Ox1R Signaling in IntA-Induced Addiction Behaviors

We found here that Ox1R signaling is critical for the maintenance of the IntA-induced ‘addiction-like’ state. We previously reported that the Ox1R antagonist SB reversed IntA-induced increases in motivation (decreases in a) and cue-induced reinstatement for cocaine at lower doses than was required for similar effects in LgA or ShA rats^5^. Here, we show that SB reversed IntA-induced increases in motivation (decreases in a) for fentanyl as well as in cue-induced reinstatement, and that SB reduced these behaviors at lower doses compared to ShA rats. We also found that the efficacy of SB in attenuating these behaviors is greatest in IntA rats. Although the mechanism for this change is unknown, increased sensitivity to Ox1R antagonism may reflect increased OxR1 signaling after IntA. This could result from increased orexin peptide release (consistent with the increased numbers of orexin-expressing neurons observed here for IntA animals), or from altered Ox1R expression in brain regions associated with drug reward such as ventral tegmental area (VTA), paraventricular thalamus (PVT) or ventral palladium (VP)^15,38–46^. One important question not addressed here is the potential role of Ox2R signaling. Previous studies demonstrate a limited role for Ox2R signaling in drug self-administration^14,47–49^. However, there is emerging evidence to indicate that Ox2Rs may contribute to addiction behaviors during enhanced motivational states^17,49^.

Importantly, the effects of SB on fentanyl demand and reinstatement behavior are unlikely due to motor impairments. We previously reported that systemic pretreatment with SB at doses identical to those used here had no effect on nose poking for sucrose in rats concurrently receiving i.v. fentanyl, nor any effect on lever pressing for sucrose in fentanyl experienced rats^9^. This is consistent with previous studies demonstrating no effect of systemic SB on general locomotor activity in either an open field apparatus^5,12^ or a cognitive task requiring motor engagement^50^. Moreover, here we found that SB treatment had no effect on inactive lever responding during cued reinstatement tests across all treatment groups, further supporting a selective effect of SB on motivated, drug-seeking behaviors.

### Increased Numbers of Orexin Neurons After IntA to Fentanyl

Orexin neurons are highly dynamic and increased numbers of orexin neurons are observed in response to several drugs of abuse^4–6,51^. We recently reported that IntA to cocaine produces a robust increase in the number of LH, but not DMH/PF, orexin-expressing neurons that persists for at least 150d of abstinence^5^. Here we report that IntA to fentanyl was associated with a persistent (at least 90d) increase in both LH and DMH orexin-expressing neurons. Because rats were sacrificed following prolonged abstinence, it is unclear if increased orexin expression associated with IntA to fentanyl is the direct result of IntA to fentanyl or prolonged abstinence. Our findings are consistent with a recent study demonstrating increased orexin neuron expression in DMH/PF and LH of mice chronically treated with non-contingent injections of morphine^6^. While the functional significance of enhanced orexin expression is unknown, it is possible that persistent increases in orexin neurons may underlie relapse risk in individuals with opioid use disorder.

Unlike with IntA to cocaine, we found that IntA to fentanyl is associated with an increase in the numbers of DMH/PF in addition to LH orexin neurons. Several lines of evidence indicate that LH orexin neurons participate in reward and motivation, whereas orexin neurons of the DMH/PF function in stress and arousal^32,52^. Augmentation of DMH/PF orexin expression associated with IntA to fentanyl may reflect the recruitment of stress systems and activation of these systems may also produce allostatic changes that lead to elevated reward thresholds during enhanced ‘addiction-like’ states^35,53,54^.

### Conclusions

Here, we report that IntA to fentanyl produced a robust ‘addiction-like’ state characterized by several endophenotypes that recapitulate DSM-5 criteria for opioid addiction. We also showed that Ox1R receptor signaling maintained IntA-induced addiction-like behaviors, and that IntA is associated with an increase in orexin cell number. Our findings demonstrate that IntA self-administration offers a robust model of opioid addiction and that perturbation of the orexin system may underly multiple aspects of the IntA induced ‘addiction-like’ state.

## Author Contributions

JEF, MHJ, and GAJ were responsible for the conception and experimental design of this project. JEF was responsible for collecting behavioral data. JEF and MHJ contributed to data analysis and interpretation. JEF drafted the manuscript, and MHJ and GAJ provided critical edits.

## Acknowledgements

We would like to thank Veronica Behman, Romina Generali, and Victoria Stritz for their assistance in performing experiments for this study.

## References

1. Mahler SV, Moorman DE, Smith RJ, James MH, Aston-Jones G. Motivational activation: a unifying hypothesis of orexin/hypocretin function. Nat Neurosci 2014;17:1298–1303.

2. Moorman DE, James MH, Kilroy EA, Aston-Jones G. Orexin/hypocretin neuron activation is correlated with alcohol seeking and preference in a topographically specific manner. Eur J Neurosci 2016;43:710–720.

3. Pantazis C, James MH, Bentzley BS, Aston-Jones G. The number of lateral hypothalamus orexin/hypocretin neurons contributes to individual differences in cocaine demand. bioRxiv 2019;547836.

4. Lawrence AJ, Cowen MS, Yang HJ, Chen F, Oldfield B. The orexin system regulates alcohol-seeking in rats. Br J Pharmacol 2006;148:752–759.

5. James MH, Stopper CM, Zimmer BA, Koll NE, Bowrey HE, Aston-Jones G. Increased Number and Activity of a Lateral Subpopulation of Hypothalamic Orexin/Hypocretin Neurons Underlies the Expression of an Addicted State in Rats. Biol Psychiatry 2018;85:925–935.

6. Thannickal TC, John J, Shan L, et al. Opiates increase the number of hypocretin-producing cells in human and mouse brain and reverse cataplexy in a mouse model of narcolepsy. Sci Transl Med 2018;10:447.

7. Matzeu A, Martin-Fardon R. Targeting the orexin system for prescription opioid use disorder: Orexin-1 receptor blockade prevents oxycodone taking and seeking in rats. Neuropharmacology 2020;164:107906.

8. Bentzley BS, Aston-Jones G. Orexin-1 receptor signaling increases motivation for cocaine-associated cues. Eur J Neurosci 2015;41:1149–1156.

9. Fragale JE, Pantazis CB, James MH, Aston-Jones G. The role of orexin-1 receptor signaling in demand for the opioid fentanyl. Neuropsychopharmacology 2019;44:1690–1697.

10. Moorman DE, Aston-Jones G. Orexin-1 receptor antagonism decreases ethanol consumption and preference selectively in high-ethanol--preferring Sprague--Dawley rats. Alcohol 2009;43:379–386.

11. Moorman DE, James MH, Kilroy EA, Aston-Jones G. Orexin/hypocretin-1 receptor antagonism reduces ethanol self-administration and reinstatement selectively in highly-motivated rats. Brain Res 2017;1654:34–42.

12. Porter-Stransky KA, Bentzley BS, Aston-Jones G. Individual differences in orexin-I receptor modulation of motivation for the opioid remifentanil. Addict Biol 2017;22:303–317.

13. Smith RJ, Aston-Jones G. Orexin / hypocretin 1 receptor antagonist reduces heroin self-administration and cue-induced heroin seeking. Eur J Neurosci 2012;35:798–804.

14. Smith RJ, See RE, Aston-Jones G. Orexin/hypocretin signaling at the orexin 1 receptor regulates cue-elicited cocaine-seeking. Eur J Neurosci 2009;30:493–503.

15. Mohammadkhani A, Fragale JE, Pantazis CB, Bowrey HE, James MH, Aston-Jones G. Orexin-1 receptor signaling in ventral pallidum regulates motivation for the opioid remifentanil. J Neurosci 2019;39:9831–9837.

16. James MH, Bowrey HE, Stopper CM, Aston-Jones G. Demand elasticity predicts addiction endophenotypes and the therapeutic efficacy of an orexin/hypocretin-1 receptor antagonist in rats. Eur J Neurosci 2018;50:2602–2612.

17. Schmeichel BE, Herman MA, Roberto M, Koob GF. Hypocretin Neurotransmission Within the Central Amygdala Mediates Escalated Cocaine Self-administration and Stress-Induced Reinstatement in Rats. Biol Psychiatry 2017;81:606–615.

18. Allain F, Minogianis EA, Roberts DC, Samaha AN. How fast and how often: The pharmacokinetics of drug use are decisive in addiction. Neurosci Biobehav Rev 2015;56:166–179.

19. Beveridge TJR WP, Brewer A, Shapiro B, Mahoney JJ, Newton TF. Analyzing Human Cocaine Use Patterns to Inform Animal Addiction Model Development. College on Problems of Drug Dependence Annual Meeting; 2012; Palm Springs, CA.

20. Kawa AB, Bentzley BS, Robinson TE. Less is more: prolonged intermittent access cocaine self-administration produces incentive-sensitization and addiction-like behavior. Psychopharmacology (Berl) 2016;233:3587–3602.

21. Zimmer BA, Oleson EB, Roberts DC. The motivation to self-administer is increased after a history of spiking brain levels of cocaine. Neuropsychopharmacology 2012;37:1901–1910.

22. Allain F, Samaha AN. Revisiting long-access versus short-access cocaine self-administration in rats: intermittent intake promotes addiction symptoms independent of session length. Addict Biol 2018;24:641–651.

23. Singer BF, Fadanelli M, Kawa AB, Robinson TE. Are Cocaine-Seeking “Habits” Necessary for the Development of Addiction-Like Behavior in Rats? J Neurosci 2018;38:60–73.

24. Calipari ES, Siciliano CA, Zimmer BA, Jones SR. Brief intermittent cocaine self-administration and abstinence sensitizes cocaine effects on the dopamine transporter and increases drug seeking. Neuropsychopharmacology 2015;40:728–735.

25. O’Neal TJ, Nooney MN, Thien K, Ferguson SM. Chemogenetic modulation of accumbens direct or indirect pathways bidirectionally alters reinstatement of heroin-seeking in high-but not low-risk rats. Neuropsychopharmacology 2019.

26. Ahmed SH, Koob GF. Transition from moderate to excessive drug intake: change in hedonic set point. Science 1998;282:298–300.

27. Hursh SR, Silberberg A. Economic demand and essential value. Psychol Rev 2008;115:186–198.

28. Bentzley BS, Fender KM, Aston-Jones G. The behavioral economics of drug self-administration: a review and new analytical approach for within-session procedures. Psychopharmacology (Berl) 2013;226:113–125.

29. Bentzley BS, Jhou TC, Aston-Jones G. Economic demand predicts addiction-like behavior and therapeutic efficacy of oxytocin in the rat. Proc Natl Acad Sci U S A 2014;111:11822–11827.

30. Mahler SV, Aston-Jones GS. Fos activation of selective afferents to ventral tegmental area during cue-induced reinstatement of cocaine seeking in rats. J Neurosci 2012;32:13309–13326.

31. Lopez MF, Moorman DE, Aston-Jones G, Becker HC. The highly selective orexin/hypocretin 1 receptor antagonist GSK1059865 potently reduces ethanol drinking in ethanol dependent mice. Brain Res 2016;1636:74–80.

32. Harris GC, Aston-Jones G. Arousal and reward: a dichotomy in orexin function. Trends Neurosci 2006;29:571–577.

33. Peyron C, Tighe DK, van den Pol AN, et al. Neurons containing hypocretin (orexin) project to multiple neuronal systems. J Neurosci 1998;18:9996–10015.

34. Skofitsch G, Jacobowitz DM, Zamir N. Immunohistochemical localization of a melanin concentrating hormone-like peptide in the rat brain. Brain Res Bull 1985;15:635–649.

35. Ahmed SH, Koob GF. Transition to drug addiction: a negative reinforcement model based on an allostatic decrease in reward function. Psychopharmacology (Berl) 2005;180:473–490.

36. Edwards S, Koob GF. Escalation of drug self-administration as a hallmark of persistent addiction liability. Behav Pharmacol 2013;24:356–362.

37. Wade CL, Vendruscolo LF, Schlosburg JE, Hernandez DO, Koob GF. Compulsive-like responding for opioid analgesics in rats with extended access. Neuropsychopharmacology 2015;40:421–428.

38. Harris GC, Wimmer M, Randall-Thompson JF, Aston-Jones G. Lateral hypothalamic orexin neurons are critically involved in learning to associate an environment with morphine reward. Behav Brain Res 2007;183:43–51.

39. Baimel C, Borgland SL. Orexin Signaling in the VTA Gates Morphine-Induced Synaptic Plasticity. J Neurosci 2015;35:7295–7303.

40. Farahimanesh S, Zarrabian S, Haghparast A. Role of orexin receptors in the ventral tegmental area on acquisition and expression of morphine-induced conditioned place preference in the rats. Neuropeptides 2017;66:45–51.

41. Dallimore JE, Mickiewicz AL, Napier TC. Intra-ventral pallidal glutamate antagonists block expression of morphine-induced place preference. Behavioral neuroscience 2006;120:1103–1114.

42. Hubner CB, Koob GF. The ventral pallidum plays a role in mediating cocaine and heroin self-administration in the rat. Brain Res 1990;508:20–29.

43. Espana RA, Oleson EB, Locke JL, Brookshire BR, Roberts DC, Jones SR. The hypocretin-orexin system regulates cocaine self-administration via actions on the mesolimbic dopamine system. Eur J Neurosci 2010;31:336–348.

44. Mahler SV, Smith RJ, Aston-Jones G. Interactions between VTA orexin and glutamate in cue-induced reinstatement of cocaine seeking in rats. Psychopharmacology (Berl) 2013;226:687–698.

45. Borgland SL, Taha SA, Sarti F, Fields HL, Bonci A. Orexin A in the VTA is critical for the induction of synaptic plasticity and behavioral sensitization to cocaine. Neuron 2006;49:589–601.

46. Matzeu A, Cauvi G, Kerr TM, Weiss F, Martin-Fardon R. The paraventricular nucleus of the thalamus is differentially recruited by stimuli conditioned to the availability of cocaine versus palatable food. Addict Biol 2017;22:70–77.

47. Uslaner JM, Winrow CJ, Gotter AL, et al. Selective orexin 2 receptor antagonism blocks cue-induced reinstatement, but not nicotine self-administration or nicotine-induced reinstatement. Behav Brain Res 2014;269:61–65.

48. Prince CD, Rau AR, Yorgason JT, Espana RA. Hypocretin/Orexin regulation of dopamine signaling and cocaine self-administration is mediated predominantly by hypocretin receptor 1. ACS Chem Neurosci 2015;6:138–146.

49. Schmeichel BE, Barbier E, Misra KK, et al. Hypocretin receptor 2 antagonism dose-dependently reduces escalated heroin self-administration in rats. Neuropsychopharmacology 2015;40:1123–1129.

50. Wiskerke J, James MH, Aston-Jones G. The orexin-1 receptor antagonist SB-334867 reduces motivation, but not inhibitory control, in a rat stop signal task. Brain Res 2019.

51. Collier AD, Min SS, Campbell SD, Roberts MY, Camidge K, Leibowitz SF. Maternal ethanol consumption before paternal fertilization: Stimulation of hypocretin neurogenesis and ethanol intake in zebrafish offspring. Prog Neuropsychopharmacol Biol Psychiatry 2020;96:109728.

52. James MH, Mahler SV, Moorman DE, Aston-Jones G. A Decade of Orexin/Hypocretin and Addiction: Where Are We Now? Curr Top Behav Neurosci 2017;33:247–281.

53. Koob G, Le Moal M. Addiction and the brain antireward system. Annual review of psychology 2008;59:29–53.

54. Koob GF. Neurobiology of Opioid Addiction: Opponent Process, Hyperkatifeia, and Negative Reinforcement. Biol Psychiatry 2020;87:44–53.

